# Oxidative DNA Damage Drives Apoptotic Photoreceptor Loss in *NMNAT1*-Associated Inherited Retinal Degeneration: A Therapeutic Opportunity

**DOI:** 10.1101/2025.06.05.658162

**Authors:** Hanmeng Zhang, Kevin Valestil, Eric A. Pierce

**Author notes:** To whom correspondence should be addressed at: Ocular Genomics Institute, Department of Ophthalmology, Massachusetts Eye and Ear, Harvard Medical School, Boston, MA 02114, USA.

## Abstract

Early-onset inherited retinal degenerations (IRDs), such as Leber congenital amaurosis (LCA) caused by pathogenic variants in the *NMNAT1* gene, lead to severe vision loss in children. Despite its ubiquitous expression, reduced *NMNAT1* function primarily affects photoreceptor cells (PRs) of the retina, yet the mechanisms underlying their heightened vulnerability remain incompletely understood. Here, we demonstrate that reduced NMNAT1 enzyme function due to the p.V9M mutation leads to DNA damage in PRs, characterized by the progressive accumulation of the oxidative DNA adduct 8-oxo-dG in *Nmnat1*^V9M/V9M^ mutant mice. Cells with oxidative DNA damage also demonstrate DNA double-strand breaks, as evidenced by co-staining with antibodies to phosphorylated H2AX (γH2A.X). This DNA damage correlates with apoptosis-driven PR degeneration, as evidenced by caspase-9 activation and TUNEL staining in the PRs of the *Nmnat1*^V9M/V9M^ mutant mice, while alternative cell death pathways such as necroptosis and parthanatos were not significantly activated. Treatment with the antioxidant N-acetylcysteine (NAC) effectively reduced oxidative DNA damage and retinal immune responses, mitigated apoptosis, and preserved cone PRs. Longitudinal assessment via optical coherence tomography (OCT) and electroretinography (ERG) revealed sustained structural and functional protection in NAC-treated mice. These findings establish oxidative DNA damage as a key driver of PR degeneration in the *Nmnat1*^V9M/V9M^ model and highlight NAC’s potential as a causal gene variant-independent therapeutic strategy for *NMNAT1*-associated IRD and potentially other IRDs in which oxidative DNA damage contributes to disease pathogenesis.

## Introduction

Severe early-onset inherited retinal degenerations (IRDs) are a diverse group of genetic disorders that cause progressive vision loss in children, significantly impacting their quality of life and underscoring the urgent need for effective therapies to preserve retinal function.^1,2^ This study focuses on a rare early-onset IRD caused by mutations in the *NMNAT1* gene, which encodes the nuclear protein nicotinamide mononucleotide adenylyltransferase 1, essential for the nicotinamide adenine dinucleotide (NAD^+^) biosynthesis in the nucleus.^3–6^ Although *NMNAT1* is ubiquitously expressed across tissues, mutations in this gene predominantly result in retina-specific phenotypes, most notably severe early-onset IRD (historically called Leber congenital amaurosis (LCA)) characterized by vision loss in early childhood, and is inherited in an autosomal recessive pattern. ^3–6^ Among the various *NMNAT1* mutations, one of the common ones is a substitution at the 9th codon, where valine is replaced by methionine (p.V9M).^3,7^ This mutation is associated with hallmark clinical features, including macular atrophy, nystagmus, progressive loss of retinal function, and ultimately severe vision loss.^3^ To model this condition, we developed *Nmnat1*^V9M/V9M^ mice, which recapitulate key aspects of the human disease.^8^ These mice exhibit early-onset retinal degeneration, marked by the progressive and robust loss of photoreceptors (PRs) within two months, followed by gradual degeneration of the inner retina and retinal pigment epithelium (RPE), accompanied by significantly reduced retinal function.^8^

Despite the ubiquitous expression of *NMNAT1*, the p.V9M mutation causes a reduction in NAD^+^ levels specifically within the retina, explaining the retina-specific phenotype.^9^ However, the reason(s) why PRs are the most vulnerable to loss of NMNAT1 function compared to other cells remain unclear. PRs have exceptionally high metabolic demands due to their role in phototransduction and the maintenance of the dark current, a continuous flow of ions (mainly Na⁺ and Ca²⁺) through PR outer segment in the dark, making them particularly sensitive to disruptions in cellular homeostasis.^10–13^ Nuclear NAD^+^, a crucial coenzyme, plays a central role in DNA repair, thereby maintaining cellular homeostasis and protecting genome integrity.^14–16^ Restoring NAD^+^ biosynthesis through adeno-associated virus (AAV)-mediated *NMNAT1* gene augmentation therapy has shown promise in preserving PRs, with the strongest protection achieved when *NMNAT1* is expressed specifically in PRs, highlighting the critical role of NAD^+^ in PR survival.^17^ In the *Nmnat1*^V9M/V9M^ mouse retina, we detected the activation of poly (ADP-ribose) polymerases (PARPs), DNA damage sensors critical for initiating NAD^+^-dependent DNA repair processes, specifically in the outer nuclear layer (ONL), where PRs are located.^8^ Further studies confirmed the presence of DNA damage using comet assays and localized this damage to the PR layer using γH2A.X, a marker of double-strand DNA breaks (DSB).^18^ On average, human cells experience approximately 70,000 DNA lesions per day, arising from processes such as DNA oxidation, alkylation, hydrolysis, and replication errors.^19–23^ Among these, oxidative DNA damage is a major contributor, particularly in PRs, where high metabolic activity increases the risk of DNA oxidation.^10–12,24^.

Severe and irreparable DNA damage can trigger multiple cell death pathways in PRs, including caspase-dependent apoptosis, PARP-mediated parthanatos, and RIPK1-mediated necroptosis.^25–29^ When DNA damage exceeds repair capacity, it initiates apoptotic signaling through caspase-9, a key regulator of the intrinsic apoptosis pathway.^30^ Caspase-9 triggers the downstream caspase cascade, ultimately leading to apoptotic cell death.^30^ In parallel, excessive PARP activation in response to severe DNA damage can deplete NAD^+^ levels, triggering the release of apoptosis-inducing factor (AIF) from mitochondria.^31,32^ AIF translocates to the nucleus, causing large-scale DNA fragmentation and chromatin condensation, resulting in parthanatos, a caspase-independent form of cell death. ^31,32^ Additionally, DNA damage can trigger a signaling cascade where RIPK1 interacts with RIPK3 and MLKL to form the necrosome complex, leading to necroptosis, a regulated form of necrotic cell death characterized by plasma membrane rupture, release of intracellular contents, and inflammation.^33–35^

In the *Nmnat1*^V9M/V9M^ mice, identifying the primary driver of DNA damage and the predominant cell death mechanisms in PRs is crucial for understanding the underlying mechanisms of PR degeneration and developing novel therapeutic approaches. In this study, we demonstrate that reduced NMNAT1 function in the *Nmnat1*^V9M/V9M^ mice results in the accumulation of oxidative DNA damage in PRs, marked by the accumulation of 8-oxo-dG and colocalization with γH2A.X. This damage correlates with apoptosis-driven PR degeneration, as evidenced by the sequential activation of caspase-9 and TUNEL staining in the PRs of the retina. Treatment with the antioxidant N-acetylcysteine (NAC) significantly reduced oxidative DNA damage, mitigated apoptosis, and provided sustained structural and functional protection. These findings provide valuable insights into the protective mechanisms of antioxidants in PRs and identify oxidative DNA damage as a promising therapeutic target for *NMNAT1*-associated IRD and potentially other genetic forms of retinal degeneration.

## Results

### Oxidative DNA Damage Accumulates in the PRs of *Nmnat1*^V9M/V9M^ Mutant Mice

NAD⁺ is essential for metabolic homeostasis and DNA repair, both of which are critical for maintaining the balance between DNA damage accumulation and repair.^14–16^ Using comet assays and γH2A.X immunostaining, we previously identified significantly elevated DNA damage specifically in PRs of the *Nmnat1*^V9M/V9M^ mouse retina.^18^ To determine whether oxidative DNA damage is the primary contributor to retinal degeneration, we measured the oxidative DNA lesion 8-oxo-dG, a hallmark of DNA oxidation from P14 to P27 in *Nmnat1*^V9M/V9M^ mutant mice, which spans the initial stage prior to and during PR degeneration.^36^ Our analysis revealed a progressive accumulation of 8-oxo-dG in *Nmnat1*^V9M/V9M^ retinas compared to *Nmnat1*^+/+^ controls, with the first detection at P16 and a significant increase at later time points (P18, *p* = 0.0052; P21, *p* = 0.0165; P24, *p* = 0.0069; P27, *p* = 0001. Figure 1A-B). Double staining for 8-oxo-dG and γH2A.X showed that all γH2A.X-positive cells also expressed 8-oxo-dG (Figure 1C-D), indicating that oxidative DNA damage is the primary endogenous source of DNA breaks in the retinas of *Nmnat1*^V9M/V9M^ mutant mice.^36,37^ Interestingly, a subset of 8-oxo-dG-positive cells (P21, 10.83%; P24, 21.85%; P27, 42.21%) lacked γH2A.X staining (Figure 1C, asterisk; Figure 1D), suggesting that oxidative DNA damage in these cells may not yet have led to detectable DNA strand breaks.

**Figure 1.**
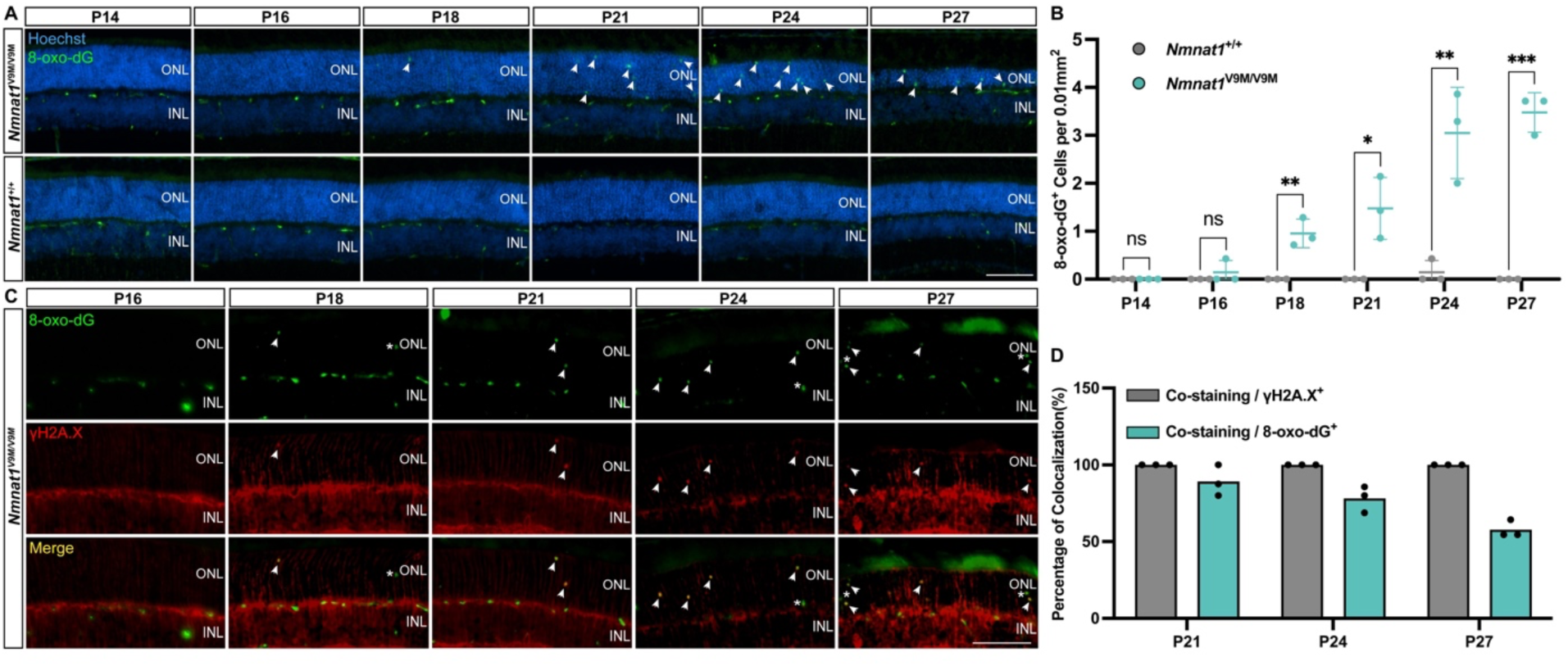
Accumulation of oxidative DNA damage in the *Nmnat1*^V9M/V9M^ mouse retina. (A) Representative immunofluorescence images showing the expression of 8-oxo-dG (green), a marker of oxidative DNA damage, in the ONL of the *Nmnat1*^V9M/V9M^ mouse retina. Nuclei are counterstained with Hoechst (blue). (B) Quantification of 8-oxo-dG levels in the ONL reveals a time-dependent increase in oxidative DNA damage in *Nmnat1*^V9M/V9M^ retinas (green) compared to *Nmnat1*^+/+^ controls (gray). Significant accumulation is observed at P18, P21, and P24. (C) Colocalization analysis of 8-oxo-dG (green) with the DNA DSB marker γH2A.X (red) in the ONL of *Nmnat1*^V9M/V9M^ retinas. Arrows indicate cells positive for both markers, while asterisks denote 8-oxo-dG-positive cells lacking γH2A.X expression. (D) Quantification of colocalization shows that all γH2A.X-positive cells (gray) also express 8-oxo-dG, while a subset of 8-oxo-dG-positive cells (green) do not colocalize with γH2A.X, suggesting oxidative DNA damage without detectable DSB formation in certain cells. Data are presented as mean ± standard deviation. Statistical comparisons between *Nmnat1*^V9M/V9M^ and *Nmnat1*^+/+^ groups were performed using multiple t-tests. N = 3 biological replicates per group. \*\*\**p* < 0.001, \*\**p* < 0.01, \**p* < 0.05; ns, non-significant. Scale bar = 100 µm.

### Oxidative DNA Damage Correlates with Apoptosis-Driven PR Degeneration in the *Nmnat1*^V9M/V9M^ Mouse Retina

To determine whether oxidative DNA damage correlates with PR degeneration, we first examined the cell death pathways activated in the retinas of *Nmnat1*^V9M/V9M^ mutant mice. Caspase-9, the initiator caspase of the intrinsic apoptotic pathway, was detected in PRs beginning at P16, followed by the late-stage apoptotic marker TUNEL at P18 (Figure 2A-B, D-E). Both caspase-9 and TUNEL signals progressively increased over time, coinciding with the progression of PR degeneration (Figure 2A-B, D-E).^30^

**Figure 2.**
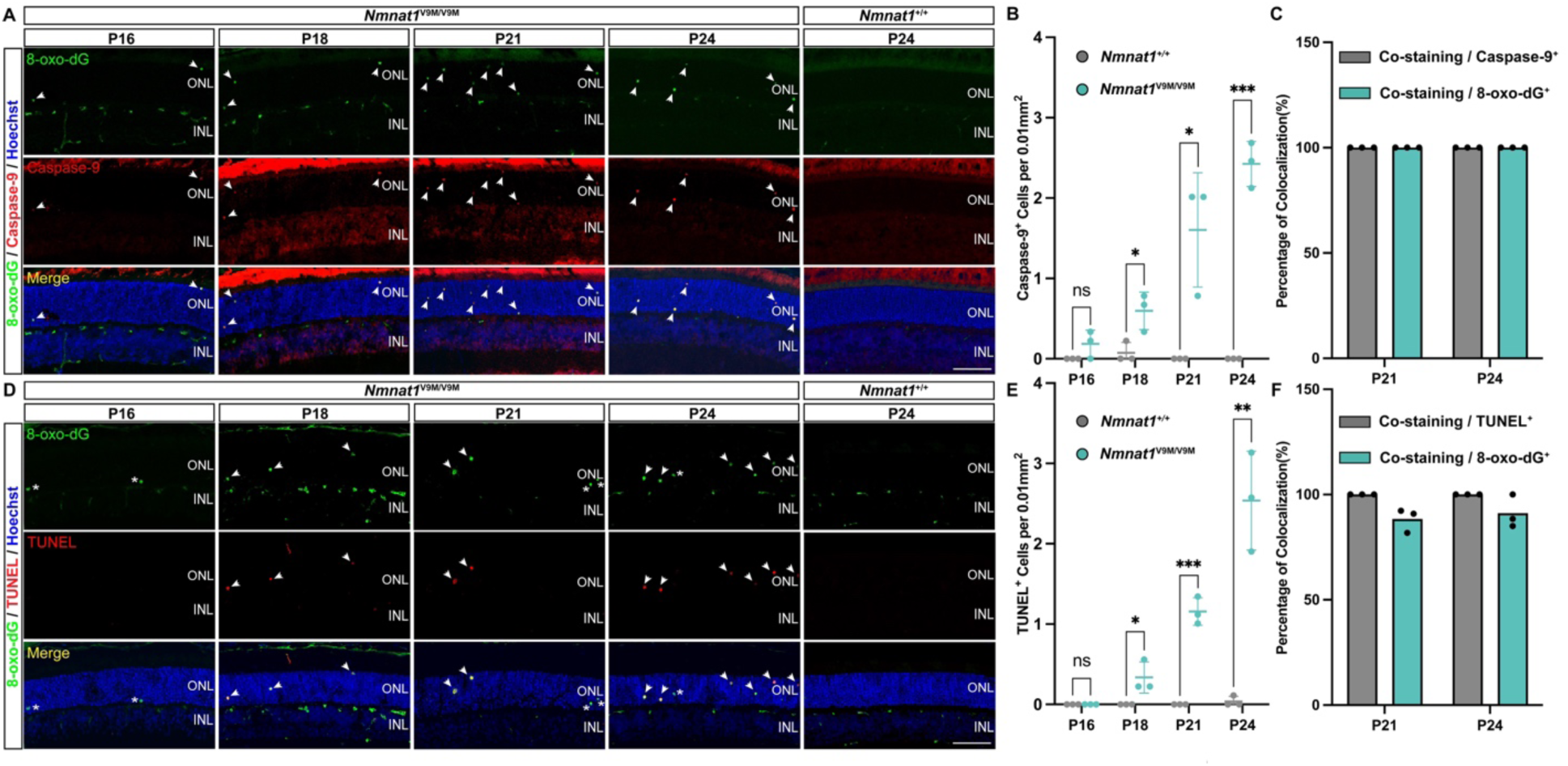
Association of oxidative DNA damage with apoptosis in the *Nmnat1*^V9M/V9M^ mouse retina. (A) Representative immunofluorescence images showing colocalization of 8-oxo-dG (green) with the apoptosis initiator caspase-9 (red) in the *Nmnat1*^V9M/V9M^ mouse retina from P16 to P24. Arrows indicate cells positive for both markers. Nuclei are counterstained with Hoechst (blue). (B) Quantification of caspase-9-positive cells reveals a time-dependent increase in apoptosis in the *Nmnat1*^V9M/V9M^ mouse retina (green) compared to *Nmnat1*^+/+^ controls (gray), with significant elevations at P18, P21, and P24. (C) Colocalization analysis confirms that all caspase-9 signals overlap with 8-oxo-dG, indicating a strong association between oxidative DNA damage and early apoptotic activation. (D) Representative images showing colocalization of 8-oxo-dG (green) with the late-stage apoptosis marker TUNEL (red) in the *Nmnat1*^V9M/V9M^ mouse retina from P16 to P24. Arrows highlight cells positive for both markers, while asterisks denote 8-oxo-dG-positive cells lacking TUNEL expression. (E) Quantification of TUNEL-positive cells demonstrates a progressive increase in apoptosis in the *Nmnat1*^V9M/V9M^ mouse retina (green) compared to controls (gray), with significant differences at P18, P21, and P24. (F) Colocalization analysis reveals that all TUNEL signals overlap with 8-oxo-dG (gray), while a subset of 8-oxo-dG-positive cells (green) do not express TUNEL, suggesting that oxidative DNA damage precedes late-stage apoptosis in some cells. Data are presented as mean ± standard deviation. Statistical comparisons between *Nmnat1*^V9M/V9M^ and *Nmnat1*^+/+^ groups were performed using multiple t-tests. N = 3 biological replicates per group. \*\*\**p* < 0.001, \*\**p* < 0.01, \**p* < 0.05; ns, non-significant. Scale bar = 100 µm.

To assess the potential involvement of alternative DNA damage-associated cell death pathways, we analyzed markers of necroptosis and parthanatos.^27–29^ Phospho-MLKL (S345), the key executioner of necroptosis, was not detected in the retinas of *Nmnat1*^V9M/V9M^ mutant mice (Supplemental Figure 1A).^28,29^ Additionally, while AIF expression was observed at P24 and increased further by P27 (Supplemental Figure 1B), no mitochondrial-to-nuclear translocation was detected (Supplemental Figure 1B), suggesting a cellular stress response rather than active parthanatos.^31,32^ Collectively, these findings indicate that apoptosis is the predominant mechanism driving PR degeneration in the retinas of *Nmnat1*^V9M/V9M^ mutant mice.

To further investigate the relationship between oxidative DNA damage and apoptotic cell death, we performed double staining for the oxidative DNA damage marker 8-oxo-dG alongside apoptotic markers from P16 to P24 (Figure 2A-B, D-E). While a subset of 8-oxo-dG-positive cells lacked TUNEL staining (Figure 2D, asterisk; Figure 2F), suggesting the possibility of DNA repair before progression to apoptosis, caspase-9 and TUNEL signals consistently colocalized with 8-oxo-dG at all time points (Figure 2A, C, D, F). These findings establish a strong correlation between oxidative DNA damage and apoptosis, reinforcing the role of oxidative DNA damage in PR degeneration in the *Nmnat1*^V9M/V9M^ mouse model.

### NAC Treatment Reduces Oxidative DNA Damage, Mitigates Apoptosis, and Preserves Cone PRs in the Retinas of *Nmnat1*^V9M/V9M^ Mutant Mice

After identifying oxidative DNA damage as the primary form of DNA damage in the retinas of *Nmnat1*^V9M/V9M^ mutant mice, we next investigated whether antioxidant treatment could mitigate this damage. To test this, we employed N-acetylcysteine (NAC), a well-established antioxidant in retinal degeneration research.^38–43^ NAC treatment was initiated at postnatal week 2 (PW2), slightly earlier than the observed significant increase in 8-oxo-dG levels around P18, ensuring antioxidant activity prior to the onset of oxidative DNA damage accumulation (Figure 1). The effects of daily NAC administration were evaluated at P18, P21, and P24, using PBS-treated *Nmnat1*^V9M/V9M^ mice as controls (Figure 3A). NAC treatment significantly reduced the number of 8-oxo-dG-positive cells in the retinas of *Nmnat1*^V9M/V9M^ mutant mice at all examined time points (P18: 89.47%, *p* = 0.0039; P21: 86.08%, *p* = 0.0006; P24: 86.76%, *p* = 0.0002; Figure 3B-C), demonstrating that antioxidant NAC effectively mitigates oxidative DNA damage accumulation.

**Figure 3.**
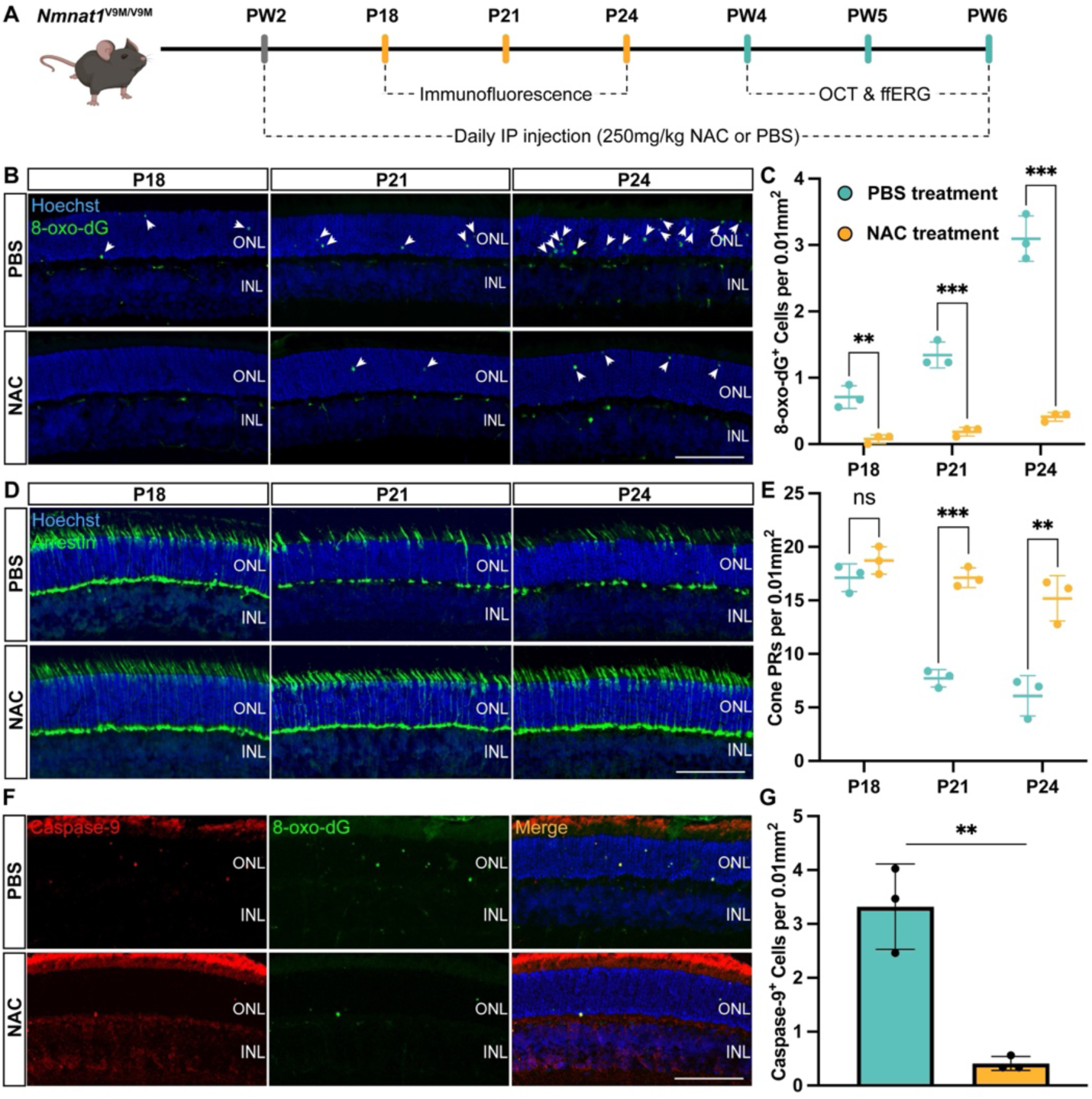
NAC treatment reduces oxidative DNA damage, mitigates apoptosis, and preserves cone photoreceptors in the Nmnat1V9M/V9M mouse retina. (A) Flowchart of the experimental design for NAC treatment. NAC administration began at PW2 via daily intraperitoneal injections. Immunofluorescence analyses were conducted from postnatal days P18 to P24, while retinal structure and functional assays were performed from PW4 to PW6. (B) Representative immunofluorescence images showing reduced 8-oxo-dG signals (green, arrows) in the NAC-treated group compared to the PBS control. Nuclei are counterstained with Hoechst (blue). (C) Quantification of 8-oxo-dG levels demonstrates that NAC treatment significantly reduces oxidative DNA damage in the *Nmnat1*^V9M/V9M^ mouse retina at P18, P21, and P24 (orange) compared to the PBS-treated control (green). (D) Representative images showing increased cone arrestin signals (red) in the NAC-treated group, indicating preserved cone PRs. (E) Quantification of cone arrestin-positive cells reveals a significant increase in the NAC-treated *Nmnat1*^V9M/V9M^ mouse retina at P21 and P24 (green) compared to the PBS-treated control (orange). (F) Representative immunofluorescence images showing reduced caspase-9 signals (red) in the NAC-treated group at P21, with colocalization to 8-oxo-dG (green), suggesting mitigation of apoptosis. (G) Quantification confirms a significant reduction in caspase-9-positive cells following NAC treatment. Data are presented as mean ± standard deviation. Statistical comparisons between NAC-treated and PBS-treated *Nmnat1*^V9M/V9M^ groups were performed using multiple t-tests. N = 3 biological replicates per group. \*\*\**p* < 0.001, \*\**p* < 0.01, \**p* < 0.05; ns, non-significant. Scale bar = 100 µm.

Given that *Nmnat1*^V9M/V9M^ mice exhibit early cone PR loss, mirroring the clinical presentation of IRD caused by *NMNAT1* mutations, we next sought to determine whether reducing oxidative DNA damage with NAC treatment could decrease cone PR death during the initial stages of disease progression.^18^ To address this, we performed double staining for caspase-9 and 8-oxo-dG at P21 following NAC treatment and assessed cone PR viability at early time points using cone arrestin as a marker. Our findings revealed that the reduction in oxidative DNA damage was accompanied by a decrease in caspase-9 levels (87.65%, *p*=0.0033; Figure 3F-G), further supporting the notion that oxidative DNA damage serves as a key upstream driver of apoptosis in the retinas of *Nmnat1*^V9M/V9M^ mutant mice. Additionally, NAC-treated *Nmnat1*^V9M/V9M^ mice exhibited significant cone PR preservation at all examined time points, including P18 (1.09-fold, *p* = 0.2022), P21 (2.22-fold, *p* = 0.0002), and P24 (2.5-fold, *p* = 0.0051) (Figure 3D-E), suggesting that early intervention to reduce oxidative DNA damage can mitigate cone PR degeneration and potentially delay more severe retinal pathology in the *Nmnat1*^V9M/V9M^ mouse retina.

### NAC Treatment Provides Sustained Protective Effects on Retinal Structure and Function

Our prior findings revealed that *Nmnat1*^V9M/V9M^ mice exhibit a rapid decline in PR layer thickness beginning at PW4.^8,18^ In addition to assessing the early therapeutic effects of NAC to investigate disease mechanisms, we next examined whether prolonged NAC administration to PW6 could confer sustained structural and functional protection, through *in vivo* optical coherence tomography (OCT) and full-field electroretinography (ffERG) analyses. Accordingly, NAC treatment was extended to PW6, and *in vivo* analyses using optical coherence tomography (OCT) and full-field electroretinography (ffERG) were performed at PW4, PW5, and PW6 (Figure 3A).

OCT imaging revealed a significant and consistent increase in outer retinal thickness (measured from the inner edge of outer plexiform layer (OPL) to the outer edge of retinal pigment epithelium (RPE)) in NAC-treated *Nmnat1*^V9M/V9M^ mice compared to PBS-treated controls. This improvement was observed across the central to peripheral retina (-600 µm to +600 µm relative to the optic nerve head center) at all time points: 1.14-to 1.25-fold at PW4 (*p* = 0.1916 to 0.0045), 1.08-to 1.33-fold at PW5 (*p* = 0.5108 to 0.015), and 1.35-to 1.48-fold at PW6 (*p* = 0.0289 to 0.0015) (Figure 4A-B). Although NAC treatment significantly preserved the outer retina, it did not fully restore retinal thickness to the levels observed in *Nmnat1*^+/+^ controls, indicating that while NAC treatment effectively slows PR degeneration and provides sustained protection, it cannot completely reverse the pathological changes caused by disrupted NMNAT1 function (Figure 4A-B). In contrast to the outer retina, inner retinal thickness (measured from the inner edge of nerve fiber layer (NFL) to the outer edge of inner nuclear layer (INL)) showed no significant differences among *Nmnat1*^+/+^, PBS-treated *Nmnat1*^V9M/V9M^, and NAC-treated *Nmnat1*^V9M/V9M^ groups (Figure 4A, C). This finding confirms that the *Nmnat1-*V9M mutation primarily affects PRs, with minimal impact on inner retinal neurons such as ganglion cells, amacrine cells, and bipolar cells at the early stages of the disease.^9,17,18,44^

**Figure 4.**
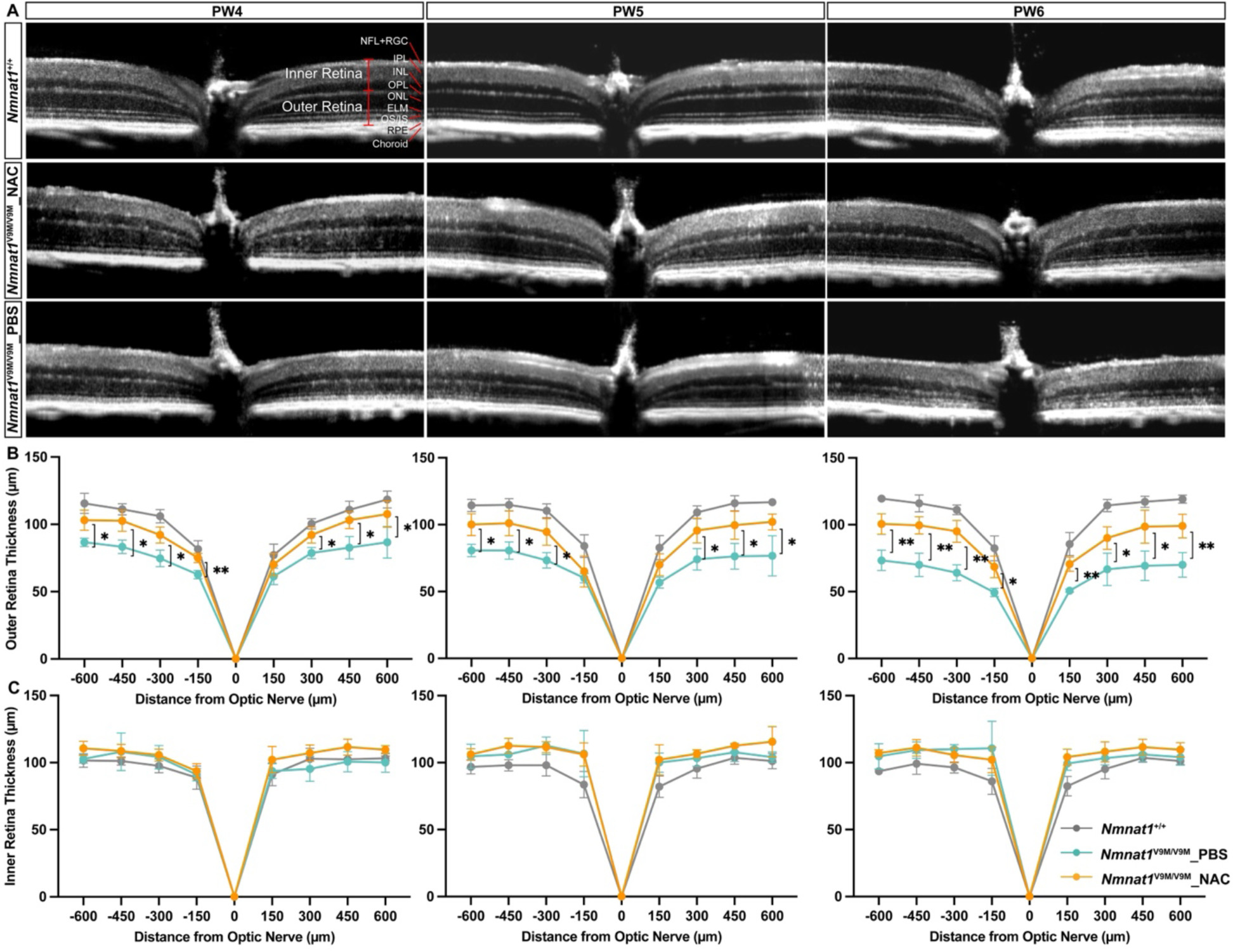
Retinal structure of NAC-and PBS-treated *Nmnat1*^V9M/V9M^ mouse retina. (A) Representative OCT scans from one eye of untreated *Nmnat1*^+/+^ (top panel), NAC-treated *Nmnat1*^V9M/V9M^ (middle panel), and PBS-treated *Nmnat1*^V9M/V9M^ mice (bottom panel) at PW4, PW5, and PW6. (B) Quantification of outer retinal thickness shows that NAC-treated *Nmnat1*^V9M/V9M^ mice (green) exhibit significant thickening compared to PBS-treated *Nmnat1*^V9M/V9M^ mice (orange) at PW4 (left panel), PW5 (middle panel), and PW6 (right panel). However, the outer retinal thickness in NAC-treated mice does not fully recover to the level of untreated *Nmnat1*^+/+^ mice (gray). (C) No significant changes were observed in inner retinal thickness among untreated *Nmnat1*^+/+^ (gray), NAC-treated *Nmnat1*^V9M/V9M^ (green), and PBS-treated *Nmnat1*^V9M/V9M^ mice (orange) at PW4 (left panel), PW5 (middle panel), and PW6 (right panel). *NFL*, nerve fiber layer; *RGC*, retinal ganglion cell; *IPL*, inner plexiform layer; *INL*, inner nuclear layer; *OPL*, outer plexiform layer; *ONL*, outer nuclear layer; *ELM*, external limiting membrane; *OS*, outer segment; *IS*, inner segment; *RPE*, retinal pigment epithelium. The inner retina includes layers from the inner edge of the NFL to the outer edge of the INL, while the outer retina includes layers from the inner edge of the OPL to the outer edge of the RPE. Data are presented as mean ± standard deviation. Statistical comparisons between NAC-treated and PBS-treated *Nmnat1*^V9M/V9M^ groups were performed using multiple t-tests. N = 4 to 5 mice per group. \*\*\**p* < 0.001, \*\**p* < 0.01, \**p* < 0.05

Consistent with the structural findings from OCT imaging, NAC treatment partially preserved retinal function in *Nmnat1*^V9M/V9M^ mice. The ffERG responses of NAC-treated *Nmnat1*^V9M/V9M^ mice were significantly improved compared to PBS-treated control mice at PW4, PW5, and PW6, although they remained below wild-type (WT, *Nmnat1*^+/+^) levels in most cases (Figure 5). Under scotopic (rod-dominant) and photopic (cone-dominant) single-flash ERG, NAC-treated mice exhibited significantly higher a-wave amplitudes, reflecting PR responses, and b-wave amplitudes, representing bipolar cell activity and PR-bipolar cell interactions, compared to PBS-treated *Nmnat1*^V9M/V9M^ controls (Figure 5A-B, D-F, H-J, L-N). These functional improvements were maintained over the two-week evaluation period and were pronounced at PW6, a stage when PR degeneration is typically more severe. Specifically, NAC treatment led to a 0.73- to 2.12-fold increase in scotopic a-wave amplitudes (*p* = 0.9772 to 0.0093), a 1.23-to 1.91-fold increase in scotopic b-wave amplitudes (*p* = 0.6953 to 0.0075), and a 1.9-to 2.72-fold increase in photopic b-wave amplitudes (*p* = 0.0408 to 0.0037). In photopic flicker ERG at 10 Hz, a condition optimized to suppress rod activity and isolate cone function, b-wave amplitudes were significantly increased in NAC-treated *Nmnat1*^V9M/V9M^ mice compared to PBS-treated *Nmnat1*^V9M/V9M^ mice (PW4: 1.64-fold increase at 10 Hz, *p* = 0.0871; PW5: 1.8-fold increase at 10 Hz, *p* = 0.0251; PW6: 2.37-fold increase at 10 Hz, *p* = 0.0018. Figure 5C, G, K, O), further demonstrating improved cone function with a sustained effect. These findings suggest that NAC provides a sustained protective effect by significantly reducing PR degeneration and preserving functional activity.

**Figure 5.**
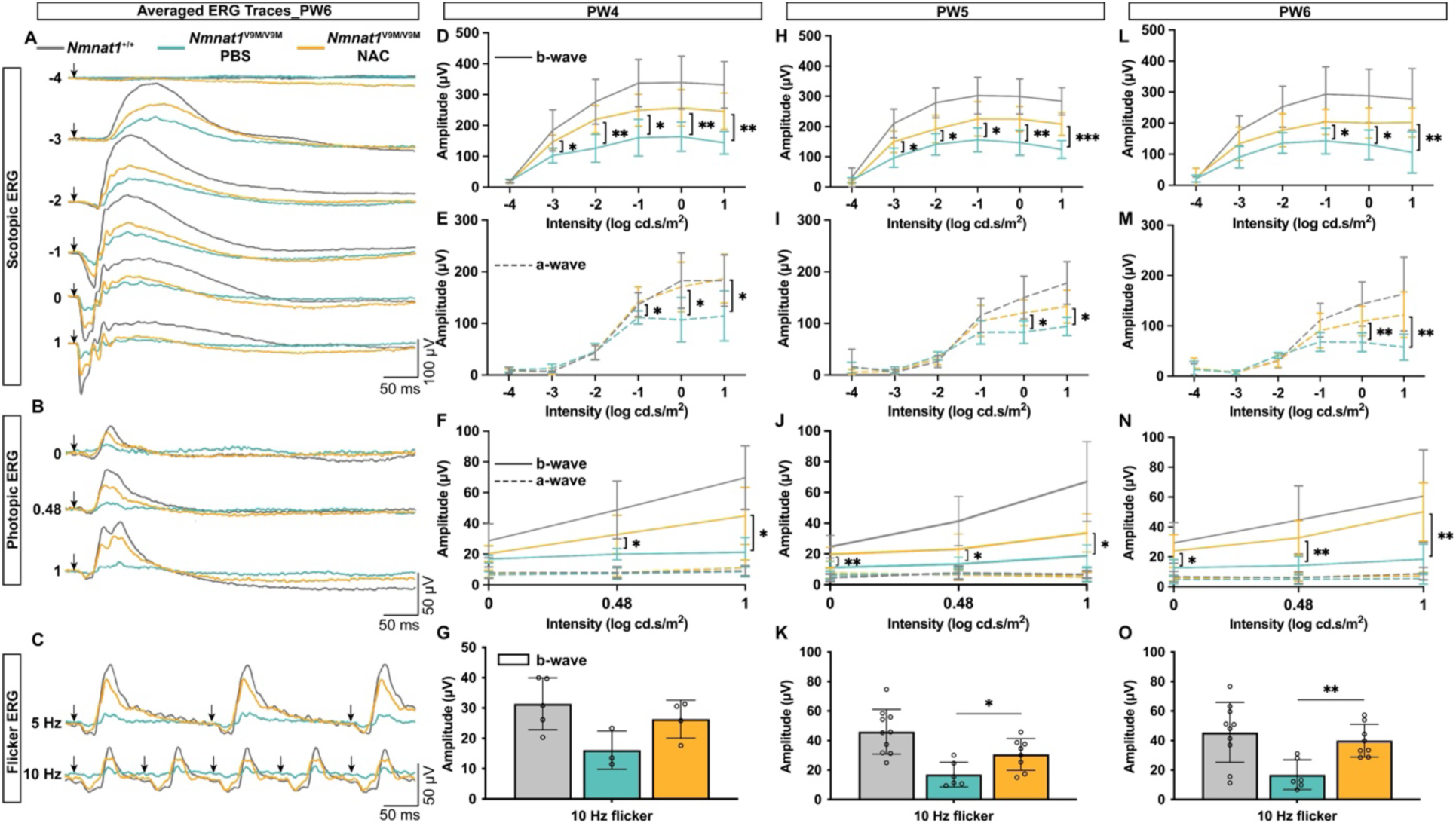
ERG analysis of NAC- and PBS-treated *Nmnat1*^V9M/V9M^ mouse retina. (A-C) Averaged ffERG traces of untreated *Nmnat1*^+/+^ (gray), NAC-treated *Nmnat1*^V9M/V9M^ (orange), and PBS-treated *Nmnat1*^V9M/V9M^ mice (green) at PW6. Arrows indicate the timing of the stimulus flash. (D-E, H-I, L-M) Statistical analysis of dark-adapted, single-flash ERG responses showing b-wave (D, H, L) and a-wave (E, I, M) amplitudes in untreated *Nmnat1*^+/+^ (gray), NAC-treated *Nmnat1*^V9M/V9M^ (orange), and PBS-treated *Nmnat1*^V9M/V9M^ mice (green) at PW4 (D-E), PW5 (H-I), and PW6 (L-M). (F, J, N) Statistical analysis of b-wave amplitudes from light-adapted, single-flash ERG responses in untreated *Nmnat1*^+/+^ (gray), NAC-treated *Nmnat1*^V9M/V9M^ (orange), and PBS-treated *Nmnat1*^V9M/V9M^ mice (green) at PW4 (F), PW5 (J), and PW6 (N). (G, K, O) Statistical analysis of b-wave amplitudes from light-adapted flicker ERG responses at 5 Hz and 10 Hz in untreated *Nmnat1*^+/+^ (gray), NAC-treated *Nmnat1*^V9M/V9M^ (orange), and PBS-treated *Nmnat1*^V9M/V9M^ (green) at PW4 (G), PW5 (K), and PW6 (O). Data are presented as mean ± standard deviation. Statistical comparisons between NAC-treated and PBS-treated *Nmnat1*^V9M/V9M^ groups were performed using multiple t-tests. N = 6 to 10 mice per group. \*\*\**p* < 0.001, \*\**p* < 0.01, \**p* < 0.05. Non-significant changes are unlabeled.

### Retinal Immune Responses Are Reduced Following NAC Treatment but Contribute Minimally to PR Degeneration in the *Nmnat1*^V9M/V9M^ Mutant Mice

In addition to its well-documented antioxidant effects, NAC has also been shown to regulate immune responses.^45,46^ Since retinal immune responses, including immune cell activation and retinal gliosis, have been observed in the retinas of *Nmnat1*^V9M/V9M^ mutant mice during early degeneration, we sought to determine whether NAC’s ability to preserve PRs also involves the modulation of retinal immune responses.^47–49^

To address this, we first characterized the time course of immune activation in the *Nmnat1*^V9M/V9M^ mouse retina from P14 to P27. Using glial fibrillary acidic protein (GFAP) as a marker for astrocytes and Müller cell processes, we observed a significant upregulation of GFAP in the *Nmnat1*^V9M/V9M^ mouse retina accompanied by the extension of GFAP-positive filaments from the inner to the outer retina starting at P18, a hallmark of gliosis (Figure 6A).^50^ Concurrently, Iba1 staining revealed reactive microglia, the resident immune cells of the retina, exhibiting hypertrophy starting at P18 (Figure 6B).^51–53^ These microglia gradually accumulated at OPL and significantly migrated toward ONL by P21 (P21, *p* = 0.0065; P24, *p* = 0.0022; P27, *p* = 0.0016. Figure 6B-C). These findings suggest that the initiation of retinal immune responses coincided with the onset of oxidative DNA damage at P18, whereas the immune cell migration to the ONL occurred slightly later, around P21, likely as a response to oxidative DNA damage in the PRs.^54–56^

**Figure 6.**
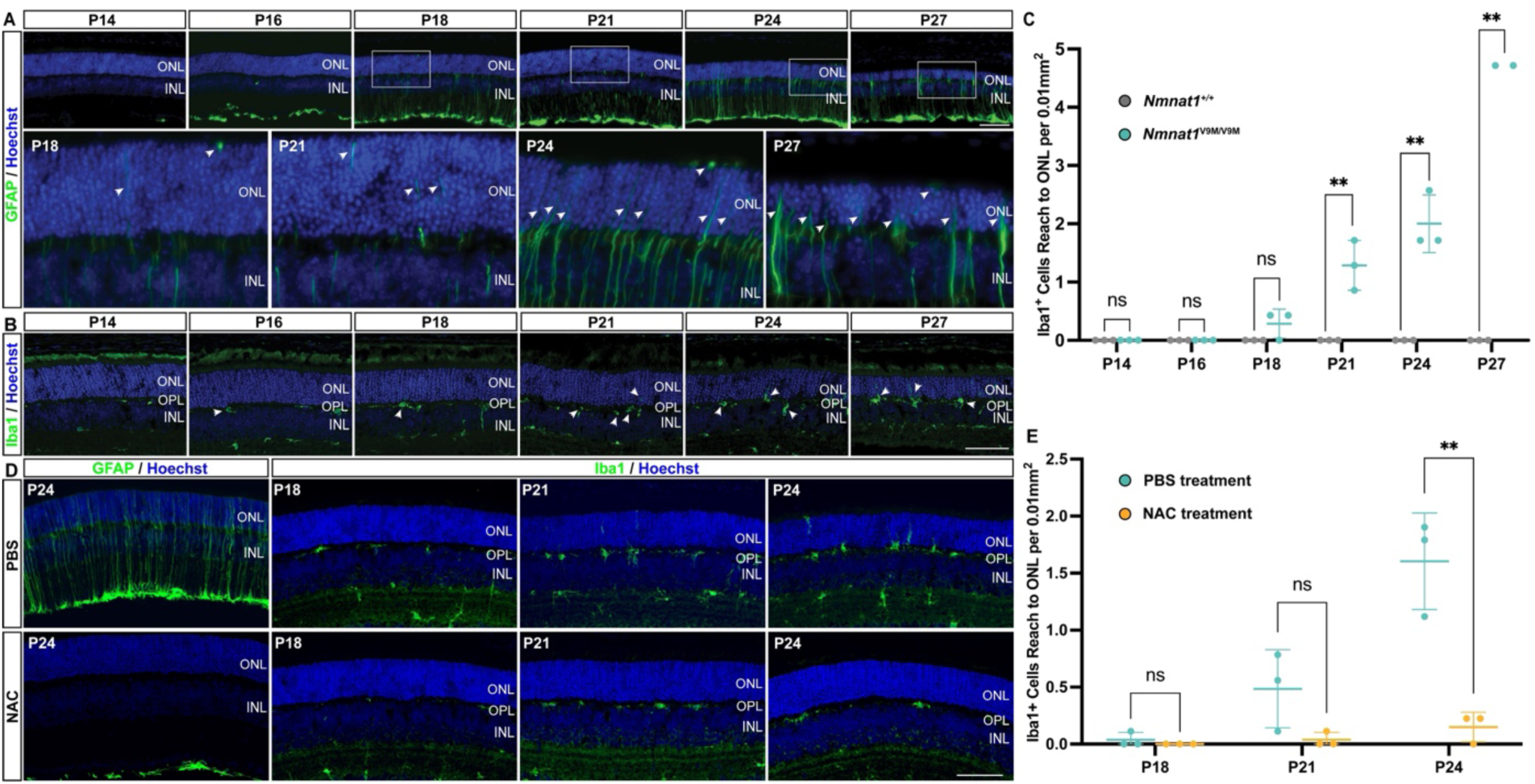
Progression of immune responses in the *Nmnat1*^V9M/V9M^ mouse retina and reduction by NAC treatment. (A) Representative images showing progressive upregulation of GFAP expression (red) in the *Nmnat1*^V9M/V9M^ retina from P14 to P27. White boxes in the top panel are magnified in the bottom panel. Arrows indicate GFAP-positive fibers extending into the ONL, with increasing intensity and distribution over time. Nuclei are counterstained with Hoechst (blue). (B) Representative images showing increased Iba1-positive microglia (green) and their migration to the ONL at later time points. Arrows highlight reactive microglia with hypertrophic morphology and their movement toward the ONL. (C) Quantification of Iba1-positive cells that have migrated to the ONL in *Nmnat1*^V9M/V9M^ (green) and *Nmnat1*^+/+^ (gray) groups, demonstrating a significant increase in microglial migration in the *Nmnat1*^V9M/V9M^ retina. (D) Representative images showing reduced GFAP overexpression and decreased Iba1-positive microglial migration to the ONL in NAC-treated *Nmnat1*^V9M/V9M^ retinas compared to PBS-treated controls. (E) Quantification of Iba1-positive cells in the ONL confirms a significant reduction in microglial migration in NAC-treated *Nmnat1*^V9M/V9M^ retinas (orange) compared to PBS-treated controls (green). Data are presented as mean ± standard deviation. Statistical comparisons between *Nmnat1*^V9M/V9M^ and *Nmnat1*^+/+^ groups or NAC-treated and PBS-treated *Nmnat1*^V9M/V9M^ groups were performed using multiple t-tests. N = 3 biological replicates per group. \*\**p* < 0.01, \**p* < 0.05; ns, non-significant. Scale bar = 100 µm.

Following NAC treatment, both GFAP-marked gliosis and Iba1-marked microglial activation and migration were significantly reduced (Figure 6D). At P24, there was a 90.7% decrease in Iba1-positive microglia reaching the ONL (*p* = 0.0047. Figure 6E). These findings suggest that NAC effectively attenuates immune responses in the *Nmnat1*^V9M/V9M^ mouse retina, either through direct immune modulation or indirectly by reducing oxidative DNA damage-induced immune activation.^47–49,54,55^

To determine whether retinal immune responses contribute significantly to PR degeneration in this mouse model, we administered a diet containing PLX5622, a colony-stimulating factor 1 receptor (CSF1R) inhibitor, to deplete microglia.^57–59^ The PLX5622 diet or a control diet was introduced at P21, coinciding with the onset of immune cell migration to the ONL (Figure 6A-C, 7A). Microglial depletion was confirmed within one week of treatment, with a significant reduction in Iba1-positive cells (88.67%, *p* = 0.0038. Figure 7B-C). However, OCT measurements of outer retinal thickness at PW4-8 showed no significant differences between PLX5622-treated and control groups (Figure 7D-E). This indicates that retinal immune responses appear to contribute minimally to PR degeneration in this model, and the protective effects of NAC on PR survival are likely mediated primarily through its reduction of oxidative DNA damage rather than modulation of immune responses.

**Figure 7.**
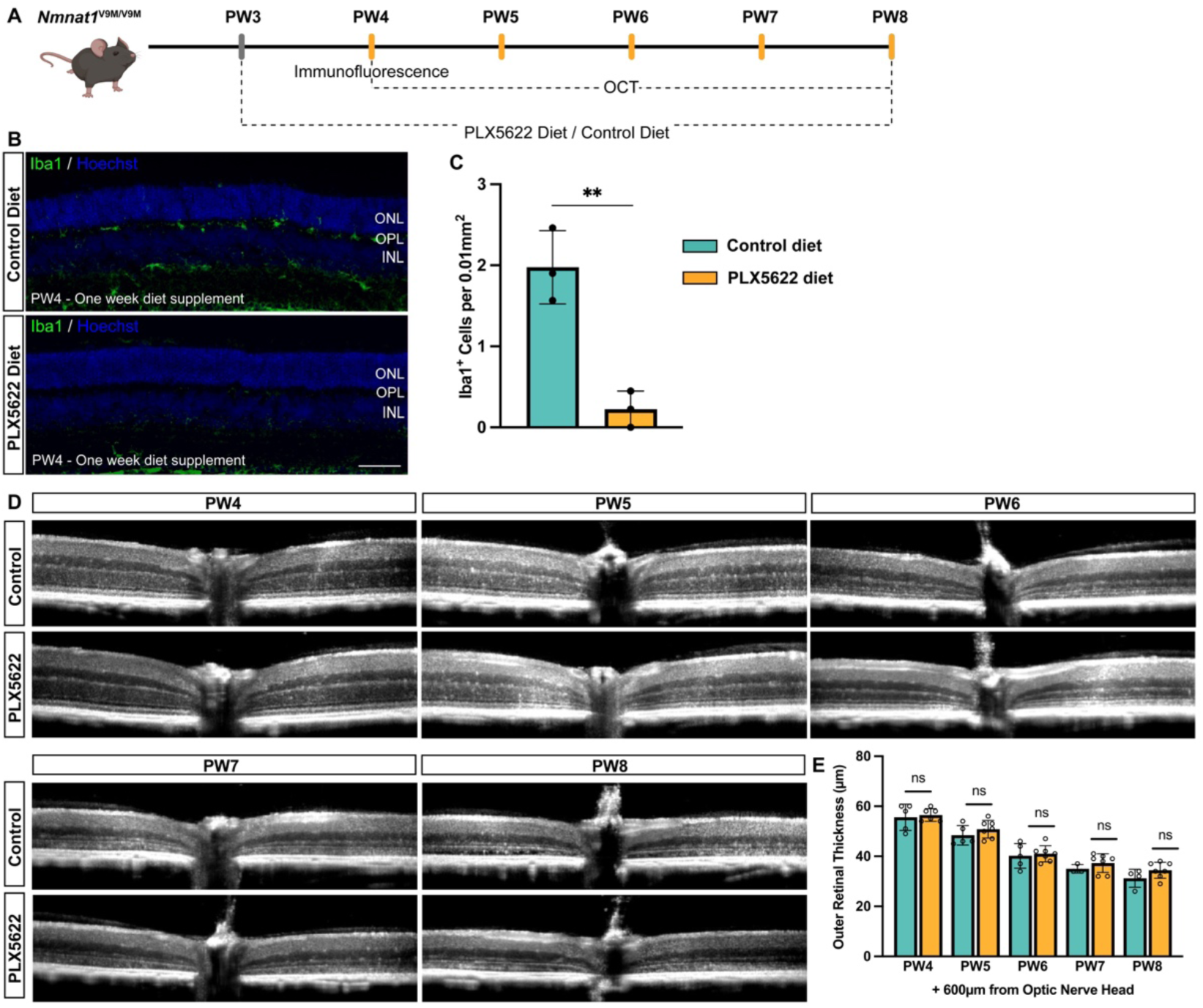
Microglia depletion does not significantly reduce PR degeneration in the *Nmnat1*^V9M/V9M^ mouse retina. (A) Flowchart of the experimental design for microglia depletion. Mice were fed a special diet containing PLX5622 or a control diet starting at PW3. Immunofluorescence was performed one week after diet initiation to assess microglia depletion efficiency. Retinal structure was evaluated *in vivo* from PW4 to PW8 to determine the protective effects of microglia depletion. (B) Representative immunofluorescence images of Iba1 (green) at one week post-diet supplementation. Markedly reduced reactive microglia were observed in the PLX5622-treated group, while control diet-treated mice showed accumulation of reactive microglia in the OPL and ONL at PW4. Nuclei are counterstained with Hoechst (blue). (C) Quantification of Iba1-positive cells reaching the ONL in PLX5622-treated (orange) and sham-treated (green) *Nmnat1*^V9M/V9M^ mouse retinas, confirming efficient microglia depletion in the PLX5622 group. (D) Representative OCT scans of PLX5622-and control diet-treated *Nmnat1*^V9M/V9M^ mouse retinas from PW4 to PW8. (E) Quantification of outer retinal thickness shows no significant difference between PLX5622-treated and sham-treated *Nmnat1*^V9M/V9M^ retinas, indicating that microglia depletion does not prevent PR degeneration. Data are presented as mean ± standard deviation. Statistical comparisons between PLX5622-treated and sham-treated *Nmnat1*^V9M/V9M^ groups were performed using multiple t-tests. N = 3 biological replicates per group for Iba1-positive cell quantification, and N = 6 mice per group for outer retinal thickness analysis. \*\**p* < 0.01; ns, non-significant. Scale bar = 100 µm.

## Discussion

Our study investigated whether oxidative DNA damage serves as a primary driver of PR degeneration in the retinas of *Nmnat1*^V9M/V9M^ mutant mice and evaluated the therapeutic potential of mitigating such damage through antioxidant intervention. The results reported demonstrated progressive accumulation of oxidative DNA damage in PRs of *Nmnat1*^V9M/V9M^ mutant mice. Further, we found that 8-oxo-dG specifically colocalized with apoptotic markers rather than markers of other cell death pathways, indicating that oxidative DNA damage causes PR cell death via apoptosis in the *Nmnat1*^V9M/V9M^ mutant mice. Of particular note, treatment with the antioxidant NAC reduced oxidative DNA damage in PRs, leading to reduced cell death and sustained protection of PR structure and function. As previously reported, retinal degeneration in the *Nmnat1*^V9M/V9M^ mutant mice was associated with a local immune response, with activation and migration of microglia into the outer retina.^18^ NAC treatment notably attenuated the retinal immune responses in the *Nmnat1*^V9M/V9M^ mice. The local immune response in the *Nmnat1*^V9M/V9M^ mice does not appear to significantly contribute to PR degeneration, as treatment with PLX5622 to deplete microglia did not alter the retinal degeneration phenotype.^57–59^ Overall, these results highlight oxidative DNA damage as a promising therapeutic target for *Nmnat1*-associated retinal degeneration and possibly other retinal degenerations sharing similar pathological mechanisms. The protective effects of NAC in the *Nmnat1*^V9M/V9M^ mouse model further strengthen the case for its therapeutic potential in PR degeneration, supporting investigations into its clinical translation.^41,60^

In our study, we use 8-oxo-dG as a reliable marker of oxidative DNA damage severity, as guanine is the most susceptible nucleobase to oxidation in DNA due to its lowest one-electron reduction potential (+1.29 V). ^61,62^ Under normal conditions, healthy cells maintain minimal 8-oxo-dG levels due to a tightly balanced system where reactive oxygen species (ROS) generate it and DNA repair mechanisms clear it. In the *Nmnat1*^V9M/V9M^ mutant mouse retina, however, we observe the overexpression and accumulation of 8-oxo-dG (Figure 1), suggesting a disruption of this balance, likely due to impaired DNA repair, as no major metabolic alterations or elevated oxidative stress were detected in these mutants from our previous study.^9^ The product of *Nmnat1*, NAD⁺, plays a critical role in DNA repair as an essential cofactor for enzymes such as PARPs, sirtuins, and DNA ligases, and we have detected reduced NAD⁺ levels in the mutant retina.^9,63^ Although direct evidence linking reduced NAD⁺ to impaired DNA repair in the retina is limited, studies in other tissues, such as skin and brain, do support this connection.^64,65^ Interestingly, 8-oxo-dG accumulation appears specifically localized to PRs, despite the ubiquitous expression of *Nmnat1* (Figure 1). Why are PRs particularly vulnerable? Unlike other retinal cells, PRs face amplified ROS generation due to intense oxygen consumption, light exposure, and oxidation-prone, polyunsaturated fatty acid-rich membranes – all of which heighten their reliance on robust DNA repair to counteract oxidative damage.^10,66,67^ This unique vulnerability means that any disruption in the delicate balance between DNA oxidation and repair disproportionately affects PRs. Indeed, elevated 8-oxo-dG levels in PRs are not unique to the *Nmnat1* mutation but are also observed in other genetic forms of IRD and light-induced retinal damage, suggesting that oxidative DNA damage is a common pathological hallmark across diverse forms of PR degeneration.^68,69^

Our findings reveal a strong correlation between oxidative DNA damage and apoptosis (Figure 2), indicating that apoptosis is the predominant mechanism driving PR degeneration in *Nmnat1*-associated IRD. This aligns with established evidence that DNA damage triggers apoptotic PR death, as observed in the light-induced PR degeneration model, and supported by our bulk RNA-seq data showing enrichment of apoptotic pathways.^18,25^ In particular, we detected caspase-9 activation (Figure 2A-C), a hallmark of the intrinsic apoptotic pathway typically initiated by cellular stressors such as DNA damage and oxidative stress.^30^ However, to our surprise, although our earlier work revealed PARP activation and reduced NAD⁺ levels – features typically associated with parthanatos, a PARP-dependent cell death pathway implicated in neurodegenerative diseases such as Alzheimer’s and Parkinson’s disease – this mechanism appears to have a limited role in *Nmnat1*-associated PR degeneration.^31,32,70,71^ This suggests that the threshold for parthanatos may not be strongly influenced by the original cellular NAD⁺ levels. Instead, in this context, PARPs likely function primarily as DNA damage sensors that initiate DNA repair, while insufficient to fully meet demand, it does not progress to NAD^+^ depletion required for parthanotic cell death.^72,73^

As an antioxidant, NAC functions both by directly scavenging ROS and serving as a precursor for glutathione synthesis, making it highly effective in models of oxidative stress–driven PR degeneration, such as retinitis pigmentosa and light-induced damage.^40,74^ Beyond its established role in mitigating oxidative stress, our study shows NAC treatment significantly reduced oxidative DNA damage accumulation and subsequent apoptotic cell death, resulting in improved PR survival (Figure 3), suggest NAC may help rebalance the oxidative DNA damage-repair equilibrium, potentially by lowering the baseline ROS burden and thus reducing the demand on DNA repair systems. This expanded hypothesis implies NAC could have broader therapeutic applications, potentially benefiting not only various forms of IRDs but also other disorders where oxidative DNA damage contributes to cellular degeneration. ^40,74–76^ The clinical potential of NAC is already being realized in retinitis pigmentosa, where trials have demonstrated both safety and preliminary effectiveness.^41,60^ Notably, the ongoing Phase 3 “NAC Attack” trial (NCT05537220) is evaluating its long-term effects on PR preservation in retinitis pigmentosa patients. Given its established safety profile and accessibility, our findings support considering NAC as both a standalone and adjunct therapy for *Nmnat1*-associated degeneration and other disorders, potentially complementing existing strategies, such as gene augmentation therapy.^17,41,60^

In *Nmnat1*-associated LCA, cones degenerate earlier than rods.^4,44,77^ An intriguing observation from our study is that the protective effect of NAC is more pronounced in cones, which exhibited a greater improvement in ERG responses, showing a 1.9-to 2.72-fold increase in photopic b-wave amplitudes, compared to a 1.23-to 1.91-fold increase in scotopic b-wave amplitudes (Figure 5). Although cones may have a higher baseline DNA repair capacity than rods, as suggested by their longer survival following DNA damage induced by alkylating agents, our findings suggest that DNA repair activity in cones may be more sensitive to NAD⁺ reduction.^78^ This vulnerability likely stems from the fundamental metabolic divergence between rods and cones. To adapt rapidly to changes in light, cones rely heavily on glycolysis for fast ATP production, whereas rods depend more on oxidative phosphorylation, which generates larger amounts of ATP to sustain the dark current.^79^ Among these processes, glycolysis is particularly sensitive to NAD⁺ depletion, and a reduction in NAD⁺ can impair DNA repair by depriving it of the required energy and building blocks and promoting chromatin condensation.^80,81^ Although metabolic changes were not detected in our previous whole-retina study, the small population of cones (∼2% of all retinal cells) may have obscured cell type-specific effects.^18,82^ The possibility that rods and cones have distinct responses to NMNAT1 deficiency is a matter of great interest and warrants further investigation using single-cell approaches.

While the retina is an immune-privileged tissue, immune cell activation can occur frequently as a common response to retinal damage arising from various causes.^83,84^ In our study, reducing oxidative DNA damage mitigated retinal immune responses, suggesting that oxidative DNA damage may act as a trigger for immune activation (Figure 6A–C). DNA damage can initiate immune responses through the release of damage-associated molecular patterns (DAMPs), such as cyclic GMP-AMP (cGAMP), which activate innate immune pathways including the cGAS-STING signaling cascade.^85^ In our bulk RNA-sequencing study of the *Nmnat1*^V9M/V9M^ mouse retina, we detected significant upregulation of both *Cgas* (2.2-fold) and *Tmem173* (STING, 2.8-fold), supporting the involvement of DNA damage-induced immune activation mechanisms.^18^ The effect of retinal immune activation is known to be biphasic: it can promote tissue repair and clearance of cellular debris, or alternatively contribute to neurotoxicity and the phagocytosis of living PRs.^86,87^ Correspondingly, microglial depletion has been reported to either rescue PRs or exacerbate PR loss depending on the model and timing.^59,88^ In our study, microglia depletion did not produce significant benefit or harm to PRs in the *Nmnat1*^V9M/V9M^ model (Figure 7), suggesting a balanced anti-inflammatory and pro-inflammatory role of microglia during the early stage of immune response. Given the complexity of microglial behavior, the timing of depletion is critical, ideally targeting phases when microglia are engaged in pathological phagocytosis rather than essential homeostatic functions. Alternatively, modulating microglial phenotype—shifting them from a pro-inflammatory (M1-like) state to an anti-inflammatory (M2-like) state—may represent a safer and more effective therapeutic approach.^86,87,89^ This could be paired with NAC treatment to target two key pathogenic events for enhanced neuroprotection.

Together, our findings demonstrate that oxidative DNA damage is a key driver of apoptosis in *Nmnat1*^V9M/V9M^ PR degeneration. Treatment with antioxidant NAC effectively reduces oxidative DNA damage and preserves PRs. To further elucidate the underlying mechanisms, future studies using single-cell RNA sequencing will investigate several key questions: whether DNA repair signaling pathways are disrupted in *Nmnat1*^V9M/V9M^ mouse PRs, what mechanisms drive early degeneration in cones, how oxidative DNA damage is linked to retinal immune responses, and whether common molecular factors contribute to oxidative DNA damage across different genetic forms of IRDs. A deeper understanding of these questions will help identify broader therapeutic targets applicable to *Nmnat1*^V9M/V9M^ and other IRDs, paving the way for the development of effective pan-therapeutic strategies against oxidative DNA damage-mediated PR degeneration.

## Methods and Materials

### Animal Subjects

All experimental procedures were approved by the Institutional Animal Care and Use Committee (IACUC) of Schepens Eye Research Institute, Massachusetts Eye and Ear, and adhered to the Association for Research in Vision and Ophthalmology (ARVO) guidelines for the use of animals in ophthalmic and vision research. Mice were housed under a 12-hour light/dark cycle and provided with a 4% fat rodent diet and water *ad libitum*. The *Nmnat1*^V9M/V9M^ mouse line was generated and maintained as previously described.^8^ For the tracking of oxidative DNA damage, retinal immune responses, and cell death markers, *Nmnat1*^V9M/V9M^ mice were bred to produce homozygous *Nmnat1*^V9M/V9M^ offspring. In antioxidant treatment experiments, *Nmnat1*^V9M/+^ mice were bred to produce litters containing both *Nmnat1*^V9M/V9M^ and *Nmnat1*^+/+^ siblings. Experiments were conducted using mice of both sexes without preference.

### Genotyping

Genomic DNA was extracted from toe biopsies using the DirectPCR lysis reagent (Viagen Biotech, Los Angeles, USA). The *Nmnat1* c.25G>A (p.V9M) mutation was identified via PCR amplification with primers (forward: 5′-CATGGCTGTGCTGAGGTG-3′; reverse: 5′-AACAGCCTGAGGTGCATGTT-3′) followed by Sanger sequencing (primer: 5′-ACGTATTTGCCCACCTGTCT-3′). PCR reactions were performed using Q5® High-Fidelity 2X Master Mix (New England Biolabs, Ipswich, USA) under standard thermocycling conditions (98°C for 30 s; 30 cycles of 98°C/10 s, 68°C/20 s, 72°C/20 s; final extension at 72°C/2 min).^8^

### Drug Administration

Starting at PW2, *Nmnat1*^V9M/V9M^ mice received daily intraperitoneal injections of NAC (Sigma-Aldrich, St. Louis, USA) dissolved in phosphate-buffered saline (PBS, 12.5 mg/mL) at a dose of 250 mg/kg body weight for 4 weeks. Control littermates received equivalent volumes of PBS (20 mL/kg). Both NAC and PBS solutions were passed through a 0.2µm polyethersulfone membrane filter (Thermo Fisher Scientific, Waltham, USA) to remove potential bacterial contaminants before injection. The NAC dosage was selected based on previous studies demonstrating ocular protection without toxicity.^38,90,91^

To achieve retinal microglial depletion, mice were given PLX5622-formulated (Chemgood, Henrico, USA) AIN-76A chow (1200 ppm, Research Diets, New Brunswick, USA) ad libitum starting at PW3.^57–59^ Control mice were fed standard AIN-76A chow (Research Diets, New Brunswick, USA). No noticeable behavioral abnormalities or health issues were observed in mice receiving the PLX5622-supplemented diet.

### OCT

Retinal structure was assessed in NAC-and PBS-treated *Nmnat1*^V9M/V9M^ mice, as well as untreated *Nmnat1*^+/+^ controls at PW4, 5, and 6 using a Bioptigen Envisu R2200 spectral-domain OCT system (Leica Microsystems, Wetzlar, Germany). Mice anesthesia was induced with 4% isoflurane and maintained at 1.5%–3% in 100% oxygen (0.4 L/min). Mydriasis was achieved using topical 2.5% phenylephrine (Lifestar Pharma, Mahwah, USA) and 1% tropicamide (Bausch + Lomb, Vaughan, Canada). Corneal hydration and image quality were maintained with periodic application of sterile saline (Sigma-Aldrich, St. Louis, USA). OCT imaging was performed using a linear B-scan centered on the optic nerve head with a 1.4 mm scan length, 1000 A-scans per B-scan, and 5 B-scan averaged over 20 frames. Retinal thickness was quantified in a blinded manner using ImageJ, with measurements taken at eccentricities spanning the optic nerve head and central retina, ranging from -600 µm to +600 µm relative to the optic nerve head.

### ffERG

Retinal function was evaluated using the Celeris ERG system (Diagnosys LLC, Lowell, USA), following previously described protocols.^92^ Prior to the procedure, all mice were dark-adapted overnight (< 24 hours), and all manipulations were conducted under dim red illumination to preserve scotopic sensitivity. Mice were anesthetized, and mydriasis was induced as described above. Corneal hydration was maintained throughout the experiment by periodically applying lubricating eye drops containing 0.3% sterile hypromellose (Alcon Laboratories, Fort Worth, USA). For scotopic ERG, brief single white flash stimuli were delivered over a five-log unit range (-4 to 1 log cd·s/m²) to assess rod and mixed rod-cone responses. Stimuli from -4 to -1 log cd·s/m² were averaged over 20 trials, while higher-intensity stimuli (0 and 1 log cd·s/m²) were averaged over five trials. Following scotopic recordings, mice were light-adapted for 10 minutes under a 25 cd/m² white background to saturate rod activity. Photopic ERG responses were then recorded using single 1 Hz flash stimuli at three intensities (0, 0.48, and 1 log cd·s/m²; 4 ms duration), with 20 responses averaged per stimulus level. Additionally, cone flicker ERGs were obtained at 10 Hz under light-adapted conditions, with 30 responses averaged per frequency. For visualization, raw ERG traces were replotted in Microsoft Excel. A-wave and b-wave amplitudes were manually identified using Espion E3 software (Diagnosys LLC, Lowell, USA) and the resulting amplitude values were exported into GraphPad Prism 10 (GraphPad Software, Boston, USA) for statistical analysis.

### Tissue Processing and Immunofluorescence

Tissue processing and immunofluorescence were conducted following established protocols.^93^ Mice were euthanized via CO₂ asphyxiation, and eyes were enucleated, rinsed in PBS, and fixed in 4% paraformaldehyde (PFA, v/v, Electron Microscopy Sciences, Hatfield, USA) for 4 minutes. The cornea and lens were dissected, and the posterior eyecup was post-fixed in 4% PFA overnight at 4°C. After PBS washes, tissues were cryoprotected in 30% sucrose (w/v) overnight at 4°C, embedded in optimal cutting temperature compound (Sakura Finetek, Torrance, USA), and sectioned at a thickness of 12 µm. For immunostaining, sections were rehydrated in PBS for 10 mins, permeabilized with 0.4% Triton X-100 (Sigma-Aldrich, St. Louis, USA) for 10 minutes, and blocked with 1% bovine serum albumin (Sigma-Aldrich, St. Louis, USA) and 10% normal goat serum (Invitrogen, Waltham, USA) in PBS for 45 minutes at room temperature. Primary antibodies were incubated overnight at 4°C, and secondary antibodies were applied sequentially at room temperature for 2 hours. Information of primary and secondary antibodies is provided in Supplemental Table 1.

For 8-oxo-dG detection, after permeabilization, RNA was enzymatically digested using RNase A (200 µg/mL, Thermo Fisher Scientific, Waltham, USA) and RNase T1 (50 U/mL, Thermo Fisher Scientific, Waltham, USA) in 140 mM NaCl (Sigma-Aldrich, St. Louis, USA) for 1 hour at 37°C, followed by a 4°C NaCl (140 mM) rinse for 5 mins. DNA denaturation was performed with 70 mM NaOH for 4 mins, and sections were treated with Proteinase K (20 µg/mL, Qiagen, Hilden, Germany) for 10 mins at 37°C. Residual enzymatic activity was quenched with 0.2% glycine (Sigma-Aldrich, St. Louis, USA) in PBS for 10 mins prior to blocking. Retinal images were captured using Nikon Eclipse Ti fluorescence microscope (Nikon, Melville, USA) equipped with an Andor CCD camera (Andor Technology, Belfast, Northern Ireland) or Leica TCS SP8 confocal microscope (Leica, Wetzlar, Germany) with standardized sampling areas equidistant from the optic nerve.

### TUNEL Assay

Late-stage apoptosis was identified using the TUNEL (TdT-mediated dUTP nick end labeling) assay (Thermo Fisher Scientific, Waltham, USA) as per the manufacturer’s protocol. Cryosections were treated with Proteinase K for 15 minutes at room temperature and washed in PBS. Sections were incubated with the TUNEL reaction mix for 60 minutes at 37°C, followed by PBS washes. For co-staining with 8-oxo-dG, TUNEL-labeled sections underwent subsequent immunostaining as described above. Nuclei were counterstained with Hoechst 33342 (Thermo Fisher Scientific, Waltham, USA).

### Statistics

Data were analyzed using Student’s t-tests with a significance threshold of α = 0.05. Results are presented as mean ± standard deviation. Detailed statistical parameters, including sample sizes and replicates, are provided in the figure legends. All analyses were performed using GraphPad Prism 10 (GraphPad Software, Boston, USA).

## Acknowledgements

The authors thank the Schepens Eye Research Institute Animal Facility staff for their help with animal care, and gratefully acknowledge Caitlin Keiper and Miele Macmillan for their technical assistance.

## Author Contribution Statement

E.A.P. and H.Z. designed the study and prepared the manuscript. H.Z. performed the experiments and analyzed data. K.V. contributed to mouse genotyping and OCT image analysis.

## Ethics Statement

All experimental animal procedures were approved by the Animal Care and Use Committee of the Schepens Eye Research Institute, Massachusetts Eye and Ear.

## Data Availability Statement

All data relevant to the study are included in the article or uploaded as supplementary information. Source data are available upon reasonable request.

## References

1. Sahel et al. Clinical Characteristics and Current Therapies for Inherited Retinal Degenerations. Cold Spring Harb Perspect Med 5 (2015) PMID25324231.

2. Bourne et al. Magnitude, temporal trends, and projections of the global prevalence of blindness and distance and near vision impairment: a systematic review and meta-analysis. Lancet Glob Health 5 e888–e897 (2017) PMID28779882.

3. Falk et al. NMNAT1 mutations cause Leber congenital amaurosis. Nature Genetics 2012 44:9 44 1040–1045 (2012) PMID22842227.

4. Perrault et al. Mutations in NMNAT1 cause Leber congenital amaurosis with early-onset severe macular and optic atrophy. Nat Genet 44 975–977 (2012) PMID22842229.

5. Chiang et al. Exome sequencing identifies NMNAT1 mutations as a cause of Leber congenital amaurosis. Nat Genet 44 972–974 (2012) PMID22842231.

6. Koenekoop et al. Mutations in NMNAT1 cause Leber congenital amaurosis and identify a new disease pathway for retinal degeneration. Nat Genet 44 1035–1039 (2012) PMID22842230.

7. Sasaki et al. Characterization of Leber Congenital Amaurosis-associated NMNAT1 Mutants. Journal of Biological Chemistry 290 17228–17238 (2015) PMID26018082.

8. Greenwald et al. Mouse Models of NMNAT1-Leber Congenital Amaurosis (LCA9) Recapitulate Key Features of the Human Disease. Am J Pathol 186 1925 (2016) PMID27207593.

9. Greenwald et al. Mutant Nmnat1 leads to a retina-specific decrease of NAD+ accompanied by increased poly(ADP-ribose) in a mouse model of NMNAT1-associated retinal degeneration. Hum Mol Genet 30 644–657 (2021) PMID33709122.

10. Pan et al. Photoreceptor metabolic reprogramming: current understanding and therapeutic implications. Commun Biol 4 (2021) PMID33627778.

11. Okawa et al. ATP consumption by mammalian rod photoreceptors in darkness and in light. Curr Biol 18 1917–1921 (2008) PMID19084410.

12. Birol et al. Oxygen distribution and consumption in the macaque retina. Am J Physiol Heart Circ Physiol 293 (2007) PMID17557923.

13. Hagins et al. Dark Current and Photocurrent in Retinal Rods. Biophys J 10 380 (1970) PMID5439318.

14. Covarrubias et al. NAD+ metabolism and its roles in cellular processes during ageing. Nat Rev Mol Cell Biol 22 119 (2020) PMID33353981.

15. Xie et al. NAD+ metabolism: pathophysiologic mechanisms and therapeutic potential. Signal Transduct Target Ther 5 (2020) PMID33028824.

16. Munk et al. NAD+ regulates nucleotide metabolism and genomic DNA replication. Nature Cell Biology 2023 25:12 25 1774–1786 (2023) PMID37957325.

17. Brown et al. Expression of NMNAT1 in the photoreceptors is sufficient to prevent NMNAT1-associated retinal degeneration. Mol Ther Methods Clin Dev 29 319–328 (2023) PMID37214313.

18. Brown et al. Reduced nuclear NAD+ drives DNA damage and subsequent immune activation in the retina. Hum Mol Genet 31 1370–1388 (2022) PMID34750622.

19 . Lindahl. Instability and decay of the primary structure of DNA. Nature 362 709–715 (1993) PMID8469282.

20. Lindahl et al. Repair of endogenous DNA damage. Cold Spring Harb Symp Quant Biol 65 127– 133 (2000) PMID12760027.

21. Tubbs et al. Endogenous DNA Damage as a Source of Genomic Instability in Cancer. Cell 168 644 (2017) PMID28187286.

22. Chatterjee et al. Mechanisms of DNA damage, repair and mutagenesis. Environ Mol Mutagen 58 235 (2017) PMID28485537.

23. Hakem. DNA-damage repair; the good, the bad, and the ugly. EMBO J 27 589 (2008) PMID18285820.

24. Domènech, et al. The Relevance of Oxidative Stress in the Pathogenesis and Therapy of Retinal Dystrophies. Antioxidants 9 (2020) PMID32340220.

25. Gordon et al. DNA damage and repair in light-induced photoreceptor degeneration. Investigative Opthalmology & Visual Science 43 3511–3521 (2002).

26. Greenwald et al. Parthanatos as a Cell Death Pathway Underlying Retinal Disease. Adv Exp Med Biol 1185 323–327 (2019) PMID31884632.

27. Li et al. Astragaloside A Protects Against Photoreceptor Degeneration in Part Through Suppressing Oxidative Stress and DNA Damage-Induced Necroptosis and Inflammation in the Retina. J Inflamm Res 15 2995–3020 (2022) PMID35645574.

28. Dong et al. PJ34 Protects Photoreceptors from Cell Death by Inhibiting PARP-1 Induced Parthanatos after Experimental Retinal Detachment. Curr Eye Res 46 115–121 (2021) PMID32478624.

29. Yu et al. UVA induces retinal photoreceptor cell death via receptor interacting protein 3 kinase mediated necroptosis. Cell Death Discovery 2022 8:1 81–11 (2022).

30. Shi. Apoptosome: The Cellular Engine for the Activation of Caspase-9. Structure 10 285–288 (2002) PMID12005427.

31. Andrabi et al. Mitochondrial and nuclear cross talk in cell death: parthanatos. Ann N Y Acad Sci 1147 233–241 (2008) PMID19076445.

32. Wang et al. A nuclease that mediates cell death induced by DNA damage and poly(ADP-ribose) polymerase-1. Science 354 aad6872 (2016) PMID27846469.

33. Kaczmarek et al. Necroptosis: The Release of Damage-Associated Molecular Patterns and Its Physiological Relevance. Immunity 38 209–223 (2013) PMID23438821.

34. Biton et al. NEMO and RIP1 control cell fate in response to extensive DNA damage via TNF-α feedforward signaling. Cell 145 92–103 (2011) PMID21458669.

35. Chen et al. PUMA amplifies necroptosis signaling by activating cytosolic DNA sensors. Proc Natl Acad Sci U S A 115 3930–3935 (2018) PMID29581256.

36. Hahm et al. 8-Oxoguanine: from oxidative damage to epigenetic and epitranscriptional modification. Experimental & Molecular Medicine 2022 54:10 54 1626–1642 (2022) PMID36266447.

37. Firsanov et al. H2AX phosphorylation at the sites of DNA double-strand breaks in cultivated mammalian cells and tissues. Clin Epigenetics 2 283 (2011) PMID22704343.

38. Sano et al. Differential effects of N-acetylcysteine on retinal degeneration in two mouse models of normal tension glaucoma. Cell Death Dis 10 (2019) PMID30692515.

39. Terluk et al. N-Acetyl-L-cysteine Protects Human Retinal Pigment Epithelial Cells from Oxidative Damage: Implications for Age-Related Macular Degeneration. Oxid Med Cell Longev 2019 5174957 (2019) PMID31485293.

40. Lee et al. N-Acetylcysteine promotes long-term survival of cones in a model of retinitis pigmentosa. J Cell Physiol 226 1843–1849 (2011) PMID21506115.

41. Campochiaro et al. The mechanism of cone cell death in Retinitis Pigmentosa. Prog Retin Eye Res 62 24–37 (2018) PMID28962928.

42. Atkuri et al. N-acetylcysteine - a safe antidote for cysteine/glutathione deficiency. Curr Opin Pharmacol 7 355 (2007) PMID17602868.

43. Kerksick et al. The Antioxidant Role of Glutathione and N-Acetyl-Cysteine Supplements and Exercise-Induced Oxidative Stress. J Int Soc Sports Nutr 2 38 (2005) PMID18500954.

44. Greenwald et al. Mouse models of NMNAT1-Leber congenital amaurosis recapitulate key features of the human disease. Invest Ophthalmol Vis Sci 57 2255–2255 (2016).

45. Verhasselt et al. N-Acetyl-l-Cysteine Inhibits Primary Human T Cell Responses at the Dendritic Cell Level: Association with NF-κB Inhibition. The Journal of Immunology 162 2569–2574 (1999) PMID10072497.

46. Yoshida et al. Laboratory evidence of sustained chronic inflammatory reaction in retinitis pigmentosa. Ophthalmology 120 (2013) PMID22986110.

47. Giordani et al. N-acetylcysteine inhibits the induction of an antigen-specific antibody response down-regulating CD40 and CD27 co-stimulatory molecules. Clin Exp Immunol 129 254–264 (2002) PMID12165081.

48 . Bonnaure et al. N-acetyl cysteine regulates the phosphorylation of JAK proteins following CD40-activation of human memory B cells. Mol Immunol 62 209–218 (2014) PMID25016575.

49. Sakai et al. N-Acetylcysteine Suppresses Microglial Inflammation and Induces Mortality Dose-Dependently via Tumor Necrosis Factor-α Signaling. Int J Mol Sci 24 3798 (2023) PMID36835209.

50. Chen et al. Expression of glial fibrillary acidic protein and glutamine synthetase by Müller cells after optic nerve damage and intravitreal application of brain-derived neurotrophic factor. Glia 38 115–125 (2002) PMID11948805.

51. Imai et al. A novel gene iba1 in the major histocompatibility complex class III region encoding an EF hand protein expressed in a monocytic lineage. Biochem Biophys Res Commun 224 855–862 (1996) PMID8713135.

52. Ito et al. Microglia-specific localisation of a novel calcium binding protein, Iba1. Molecular Brain Research 57 1–9 (1998) PMID9630473.

53 . Ibrahim et al. Retinal microglial activation and inflammation induced by amadori-glycated albumin in a rat model of diabetes. Diabetes 60 1122–1133 (2011) PMID21317295.

54 . Pezone et al. Inflammation and DNA damage: cause, effect or both. Nature Reviews Rheumatology 2023 19:4 19 200–211 (2023) PMID36750681.

55. Arvanitaki et al. DNA damage, inflammation and aging: Insights from mice. Frontiers in aging 3 (2022) PMID36160606.

56. Rock et al. The inflammatory response to cell death. Annu Rev Pathol 3 99 (2008) PMID18039143.

57. Huang et al. Dual extra-retinal origins of microglia in the model of retinal microglia repopulation. Cell Discovery 2018 4:1 4 1–16 (2018).

58. Okunuki et al. Retinal microglia initiate neuroinflammation in ocular autoimmunity. Proc Natl Acad Sci U S A 116 9989–9998 (2019) PMID31023885.

59. Okunuki et al. Microglia inhibit photoreceptor cell death and regulate immune cell infiltration in response to retinal detachment. Proc Natl Acad Sci U S A 115 E6264–E6273 (2018) PMID29915052.

60. Grandjean et al. Efficacy of oral long-term N-acetylcysteine in chronic bronchopulmonary disease: A meta-analysis of published double-blind, placebo-controlled clinical trials. Clin Ther 22 209– 221 (2000).

61 . Burrows et al. Oxidative Nucleobase Modifications Leading to Strand Scission. Chem Rev 98 1109–1151 (1998) PMID11848927.

62. Cadet et al. Formation and repair of oxidatively generated damage in cellular DNA. Free Radic Biol Med 107 13–34 (2017) PMID28057600.

63. Ruszkiewicz et al. Fueling genome maintenance: On the versatile roles of NAD+ in preserving DNA integrity. Journal of Biological Chemistry 298 102037 (2022) PMID35595095.

64. Fang et al. Nuclear DNA damage signalling to mitochondria in ageing. Nat Rev Mol Cell Biol 17 308–321 (2016) PMID26956196.

65. Lautrup et al. NAD+ in Brain Aging and Neurodegenerative Disorders. Cell Metab 30 630 (2019) PMID31577933.

66. Wang et al. Role of Oxidative Stress in Retinal Disease and the Early Intervention Strategies: A Review. Oxid Med Cell Longev 2022 7836828 (2022) PMID36275903.

67. Fu et al. Fatty acid oxidation and photoreceptor metabolic needs. J Lipid Res 62 (2021) PMID32094231.

68. Murakami et al. Oxidative Stress and Microglial Response in Retinitis Pigmentosa. International Journal of Molecular Sciences 2020, Vol.21, Page 7170 21 7170 (2020) PMID32998461.

69. Xu et al. Stimulation of AMPK prevents degeneration of photoreceptors and the retinal pigment epithelium. Proceedings of the National Academy of Sciences 115 10475–10480 (2018) PMID30249643.

70. Kam et al. Poly(ADP-ribose) drives pathologic α-synuclein neurodegeneration in Parkinson’s disease. Science 362 (2018) PMID30385548.

71 . Martire et al. PARP-1 involvement in neurodegeneration: A focus on Alzheimer’s and Parkinson’s diseases. Mech Ageing Dev 146–148 53–64 (2015) PMID25881554.

72. Svilar et al. Base excision repair and lesion-dependent subpathways for repair of oxidative DNA damage. Antioxid Redox Signal 14 2491–2507 (2011) PMID20649466.

73. Bauer et al. The current state of eukaryotic DNA base damage and repair. Nucleic Acids Res 43 10083–10101 (2015) PMID26519467.

74. Nakamura et al. The involvement of the oxidative stress in murine blue LED light-induced retinal damage model. Biol Pharm Bull 40 1219–1225 (2017) PMID28769003.

75. Lovell et al. Oxidative DNA damage in mild cognitive impairment and late-stage Alzheimer’s disease. Nucleic Acids Res 35 7497–7504 (2007) PMID17947327.

76. Delint-Ramirez et al. DNA damage and its links to neuronal aging and degeneration. Neuron 113 7–28 (2025) PMID39788088.

77 . Falk et al. NMNAT1 mutations cause Leber congenital amaurosis. Nat Genet 44 1040 (2012) PMID22842227.

78. Nomura-Komoike et al. Effects of different alkylating agents on photoreceptor degeneration and proliferative response of Müller glia. Sci Rep 14 1–10 (2024) PMID38167441.

79. Chen et al. Retinal metabolism displays evidence for uncoupling of glycolysis and oxidative phosphorylation via Cori-, Cahill-, and mini-Krebs-cycle. Elife 12 (2023) PMID38739438.

80. Liu et al. Glycolytic metabolism influences global chromatin structure. Oncotarget 6 4214–4225 (2015) PMID25784656.

81. Sun et al. Impact of glycolysis enzymes and metabolites in regulating DNA damage repair in tumorigenesis and therapy. Cell Communication and Signaling 2025 23:1 23 1–17 (2025) PMID39849559.

82. Hughes et al. Cell Type-Specific Epigenomic Analysis Reveals a Uniquely Closed Chromatin Architecture in Mouse Rod Photoreceptors. Sci Rep 7 (2017) PMID28256534.

83 . Stepp et al. Immune responses to injury and their links to eye disease. Transl Res 236 52 (2021) PMID34051364.

84. O’Koren et al. Microglial Function Is Distinct in Different Anatomical Locations during Retinal Homeostasis and Degeneration. Immunity 50 723 (2019) PMID30850344.

85 . Li et al. The cGAS–cGAMP–STING pathway connects DNA damage to inflammation, senescence, and cancer. J Exp Med 215 1287 (2018) PMID29622565.

86. Orihuela et al. Microglial M1/M2 polarization and metabolic states. Br J Pharmacol 173 649–665 (2016) PMID25800044.

87. Cherry et al. Neuroinflammation and M2 microglia: The good, the bad, and the inflamed. J Neuroinflammation 11 1–15 (2014) PMID24889886.

88. Zhao et al. Microglial phagocytosis of living photoreceptors contributes to inherited retinal degeneration. EMBO Mol Med 7 1179–1197 (2015) PMID26139610.

89. Guo et al. Microglia Polarization From M1 to M2 in Neurodegenerative Diseases. Front Aging Neurosci 14 815347 (2022) PMID35250543.

90. Osada et al. Neuroprotective effect of bilberry extract in a murine model of photo-stressed retina. PLoS One 12 e0178627 (2017) PMID28570634.

91. Yu et al. NADPH and NAC synergistically inhibits chronic ocular hypertension-induced neurodegeneration and neuroinflammation through regulating p38/MAPK pathway and peroxidation. Biomedicine & Pharmacotherapy 175 116711 (2024) PMID38735082.

92. Zhang et al. Electroretinogram of the cone-dominant thirteen-lined ground squirrel during Euthermia and Hibernation in comparison with the rod-dominant Brown Norway rat. Invest Ophthalmol Vis Sci 61 6–6 (2020) PMID32492111.

93. Zhang et al. Pre-retinal delivery of recombinant adeno-associated virus vector significantly improves retinal transduction efficiency. Mol Ther Methods Clin Dev 22 96–106 (2021).

