## Supplemental Figure 1 for "Oxidative DNA Damage Drives Apoptotic Photoreceptor Loss in *NMNAT1*-Associated Inherited Retinal Degeneration: A Therapeutic Opportunity"

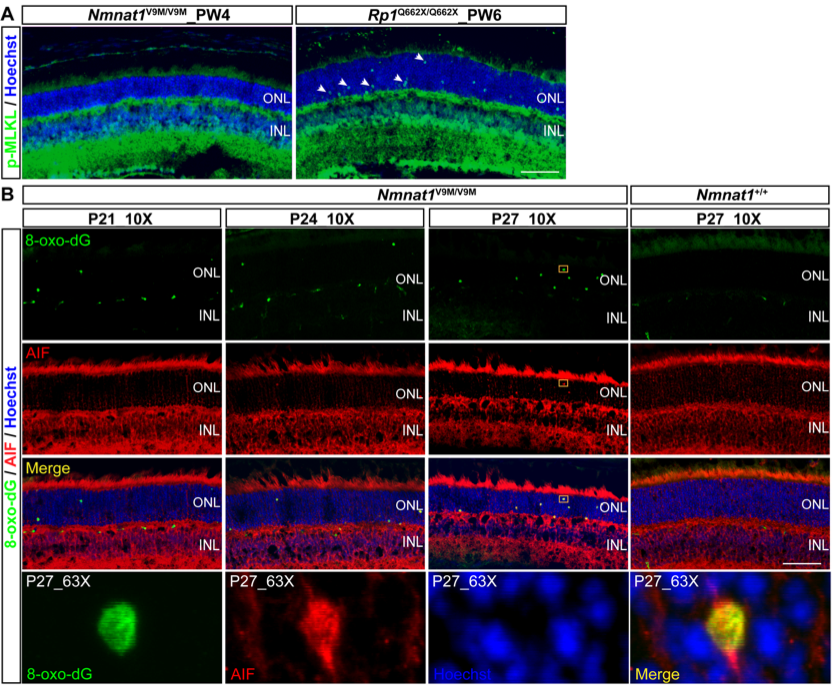

**Supplemental Figure 1. Necroptosis and parthanatos contribute minimally to *Nmnat1*-associated retinal degeneration.** (A) Immunofluorescence analysis of the necroptosis marker phospho-MLKL (green) in the *Nmnat1*^V9M/V9M^ retina at PW4 shows no detectable signal, indicating the absence of necroptosis during PR degeneration. *Rp1*^Q662X/Q662X^ mice at PW6, which undergo PR degeneration, were used as a positive control to confirm the reactivity of the phospho-MLKL antibody. Nuclei are counterstained with Hoechst (blue). (B) Immunofluorescence of AIF (red), a marker of parthanatos, reveals its presence at later time points (P24 and P27) in the *Nmnat1*^V9M/V9M^ mouse retina. Zoomed in images show that AIF colocalizes with 8-oxo-dG (green) but does not translocate to Hoechst-labeled cell nuclei, suggesting that parthanatos is not a major contributor to cell death in this model. Scale bar = 100 µm.
