## Supplemental Table 1 for "Oxidative DNA Damage Drives Apoptotic Photoreceptor Loss in *NMNAT1*-Associated Inherited Retinal Degeneration: A Therapeutic Opportunity"

**Supplemental Table 1.** List of primary and secondary antibodies used in experiments.

A. **Primary Antibody**

| **Antigen** | **Host Species** | **Catalog #** | **Dilution** | **Source** |
| --- | --- | --- | --- | --- |
| 8-oxo-dG | Mouse | 4354MC050 | 1:200 | R&D Systems |
| γH2A.X | Rabbit | 4418-APC-100 | 1:200 | Bio-Techne |
| GFAP | Rabbit | AB68428 | 1:200 | Abcam |
| Iba1 | Rabbit | 019-19741 | 1:150 | Fujifilm |
| Caspase-9 | Rabbit | AB52298 | 1:200 | Abcam |
| AIF | Rabbit | AB1998 | 1:100 | Abcam |
| p-MLKL | Rabbit | AB196436 | 1:200 | Abcam |

B. **Secondary Antibody**

| **Antigen** | **Catalog #** | **Dilution** | **Source** |
| --- | --- | --- | --- |
| Goat-anti-mouse Alexa Fluor 488 | A28175 | 1:1000 | Thermo Fisher Scientific |
| Goat-anti-rabbit Alexa Fluor 555 | A21428 | 1:1000 | Thermo Fisher Scientific |
| Hoechst 33342 | H3570 | 1:1000 | Thermo Fisher Scientific |
